# Prediction of vascular aging based on smartphone acquired PPG signals

**DOI:** 10.1101/2020.05.26.116186

**Authors:** Lorenzo Dall’Olio, Nico Curti, Daniel Remondini, Yosef Safi Harb, Folkert W. Asselbergs, Gastone Castellani, Hae-Won Uh

**Author notes:** these authors contributed equally to this work (Gastone Castellani, Hae-Won Uh).

## Abstract

Photoplethysmography (PPG) measured by smartphone has the potential for a large scale, non-invasive, and easy-to-use screening tool. Vascular aging is linked to increased arterial stiffness, which can be measured by PPG. We investigate the feasibility of using PPG to predict healthy vascular aging (HVA) based on two approaches: machine learning (ML) and deep learning (DL). We performed data preprocessing including detrending, demodulating and denoising on the raw PPG signals. For ML, ridge penalized regression has been applied to 38 features extracted from PPG, whereas for DL several convolutional neural networks (CNNs) have been applied to the whole PPG signals as input. The analysis has been conducted using the crowd-sourced Heart for Heart data. The prediction performance of ML using two features (AUC of 94.7%) – the ***a*** wave of the second derivative PPG and ***tpr***, including four covariates, sex, height, weight, and smoking – was similar to that of the best performing CNN, 12-layer ResNet (AUC of 95.3%). Without having the heavy computational cost of DL, ML might be advantageous in finding potential biomarkers for HVA prediction. The whole workflow of the procedure is clearly described, and open software has been made available to facilitate replication of the results.

## Introduction

Photoplethysmography (PPG) offers a non-invasive optical measurement method and can be used for heart rate (HR) monitoring purposes. For example, by using the white light-emitting diode as light source and the phone camera as photo-detector positioned on the index fingertip, a smartphone could be used to measure the volumetric variations of blood circulation without any additional devices^1,2^. Smartphone ownership continues to grow rapidly, and mobile phone apps are increasingly used for screening, diagnosis, and monitoring of HR and rhythm disorders such as atrial fibrillation (AF)^3,4^. The benefits and harms of using mobile apps to improve cardiovascular health or screening for specific diseases have been evaluated and discussed^3,5–7^; while HR measured by smartphone apps performing PPG agrees with a validated method such as electrocardiogram (ECG) in resting sinus rhythm^6^, the US Preventive Services Task Force (USPSTF) found that the current evidence is insufficient to assess the balance of benefits and harms of screening for AF with ECG^7^.

Our motivating crowd-sourced data comes from the Heart for Heart (H4H) initiative, promoted by the Arrhythmia Alliance, the Atrial Fibrillation Association, Happitech and other partners^8,9^. The aim of H4H is to gather millions of cardiac measurements and to increase the pace of progress on AF diagnostic technology. The primary advantages of using this population cohort data are: abundance of PPG recordings in large samples (ca. 10,000) and relatively long sequences (90 seconds), and free access to raw PPG signals via Happitech app. On the downside, these PPG signals can be very noisy compared to those in clinical settings, due to the uninstructed and not monitored PPG captures – during the measurement an appropriate pressure should be maintained and the measuring point should be kept still. Analyzing crowd-sourced data, and not data from some clinical trials, hinders to directly evaluate PPG signals for prediction of AF. Moreover, validation of the results through confirmation of ECG is not possible. On the other hand, to monitor cardiovascular status such as arterial blood pressure or arterial stiffness, the non-invasive PPG technique can be very useful.

Aging is a major non-reversible risk factor for cardiovascular disease. Chronological aging plays significant role in promoting processes involved in biological aging including oxidative stress, DNA methylation, telomere shortening, as well as structural and functional changes to the vasculature of the heart^10^. Vascular aging, in particular, is characterized by a gradual change of the vascular structure and function, and increasing arterial stiffness is considered to be the hallmark of vascular aging^11–13^. Arterial stiffness can be measured by pulse wave velocity (PWV)^14^, or by the use of the PPG technique^15^. In particular, some aging indexes (AGI) can be calculated from the second derivative of the PPG (SDPPG) waveform^16,17^. In our data, additional information such as age, sex, weight, height and smoking is available. Therefore, we consider the chronological age as a proxy for vascular aging, and investigate how well the PPG signals can predict and classify healthy or unhealthy vascular aging.

Figure 1 summarizes the procedure at a quick glance. Briefly, after detrending, demodulating and denoising the PPG signals, we consider two approaches, machine learning (ML) and deep learning (DL). While ML is based on (known) extracted features from PPG, such as beat-to-beat interval, DL uses the whole signal to predict HVA, skipping the feature extraction step. The prediction performance of the final models is validated on the dataset held back from the training and testing of the model, such that AUC (area under the curve) presents an unbiased performance measure for comparing final models. Moreover, clearly documenting the whole procedure from data preprocessing, statistical analysis, to validation of the results, will enable the scientific community to improve the usage of PPG in health care, and may lead to a robust standard operating procedure. By doing so, our work contributes to promoting open science, open software (source code in Python), and open data (available on request by H4H).

**Figure 1.**
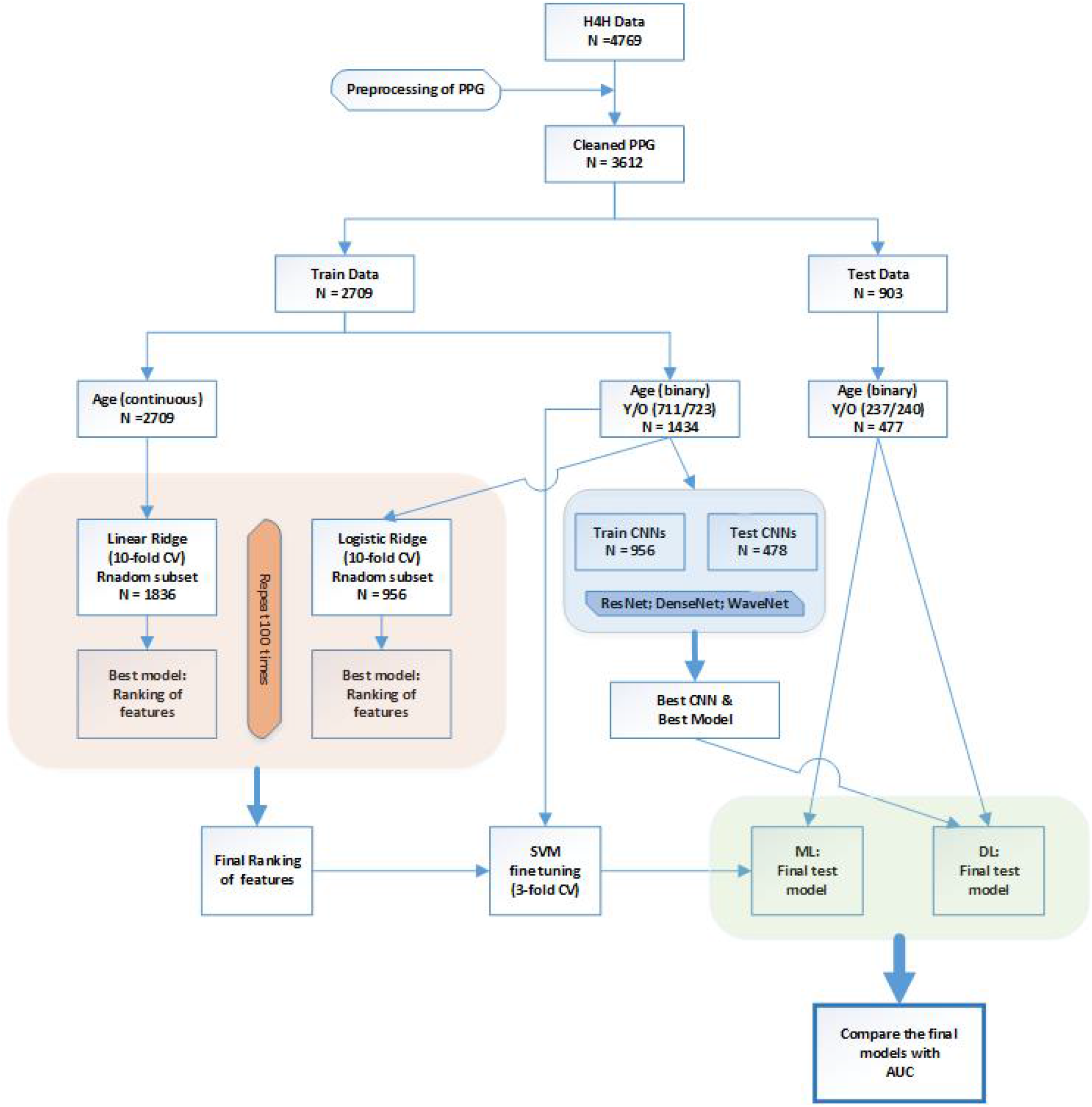
Workflow of PPG analysis to predict healthy vascular aging (HVA) using ML and DL approaches.

## Methods

### Description of datasets used

Our database comes from the Heart for Heart initiative^8^, and it consists of 4769 individuals. For each subject, one file in the .csv format is provided, containing the PPG recordings and some additional information such as sex, age, weight, height, and smoking. Data matrices of PPG recordings have 7 columns: time (around 90 seconds of measurement with a sample frequency of averagely 30 points per second), simultaneous recordings of red, green and blue light PPG, and three axis of an accelerometer. Infrared light has a more effective penetration depth in the skin compared to green light, but is more susceptible to motion artifacts^1,3,18^. For the current analysis, we assume that motion artifact in PPG signals obtained from the smartphone camera recording is negligible compared to those obtained by smartwatch. Considering the relative short recordings of PPG, we took two columns (time and red PPG) for further analysis. More detailed description is given in Supplementary Materials (Section 1 and Supplementary Fig. S1).

### PPG Preprocessing

We first removed the trend of the raw red PPG signal by computing a centered moving average (CMA) and subtracting it from the raw signal (to perform a high-pass filter). The sliding window **w** (average sampling frequency) was used to have the same number of points per second for each of the different signals. Next, to account for motion and noise artifacts, demodulation of PPG signals was obtained using the detrended signal (a real-valued *s*) and its Hilbert transform 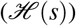 to generate the analytic signal (a complex-valued 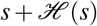). From the module of the analytic signal, the instantaneous amplitude (also called the envelope) was obtained and then smoothed with CMA. Lastly, the detrended signal was divided by the smoothed envelope, resulting in the detrended, demoduled, and denoised signal for further analysis (Fig. 2). More details can be found in Supplementary Materials (Section 2 and Supplementary Fig. S2 and S3).

**Figure 2.**
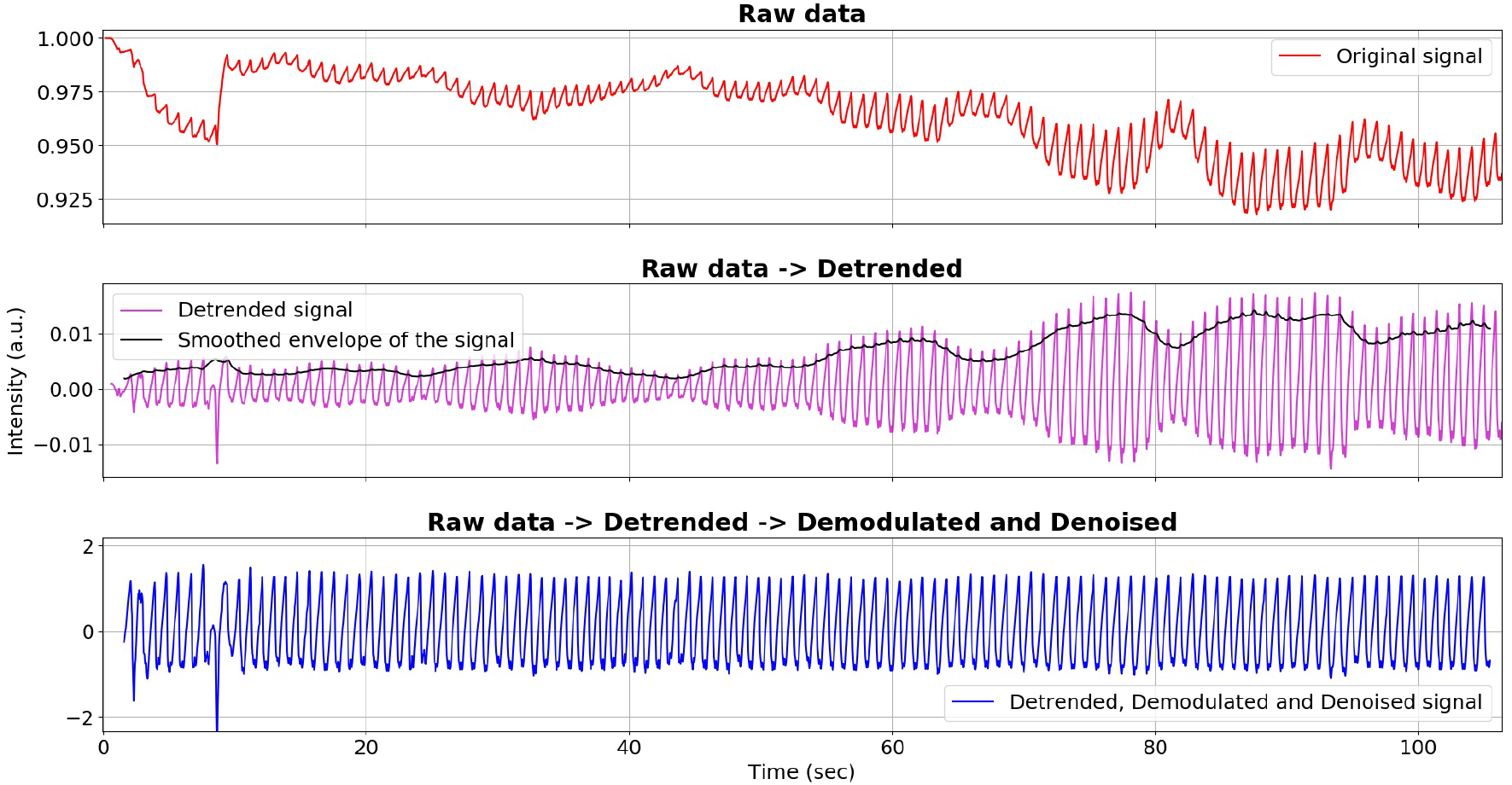
Preprocessing of PPG: (from top to bottom) raw, detrended, and demodulated and denoised clean signal.

### Feature extraction from the PPG signal for the ML approach

#### Peak detection

To identify the correct peak locations, we slightly modified an existing algorithm^19^. Our current algorithm consists of: (i) compute the CMA of the processed signal and align it to its signal; (ii) locate the maximum of a specific region of the signal continuously above its moving average; (iii) label the set of maxima as partial peaks, repeating with the window width **w**_*i*_ = (0.5, 1, 1.5, 2, 2.5, 3) ∗ *s f* (with *s f* = average sampling frequency), and consider as possible maxima those points labelled 6 times as partial peaks; (iv) if two possible maxima are separated by less than 400ms, discard the smaller of the two. Having found the final peak positions, RR can be defined as the sequence of time intervals (expressed in ms) between consecutive detected peaks.

#### Features extracted from RR

From RR we can extract information about heart rate variability (HRV) and inter beat time intervals distribution: ***ibi*** (average inter beat interval); ***medianRR***; ***madRR*** (median absolute distance); ***sdnn*** (the standard deviation); ***tpr*** (turning point ratio of RR: computed as number of local extrema divided by number of points in the signal, ranging from 0 to 1 and used as and index of randomness^20^; ***skewnessRR***; ***kurtosisRR***, etc.

#### Features extracted from RR_diff_

The features derived from *RR*_*diff*_, the sequence of differences between successive elements of RR, are the following: ***sdsd*** (standard deviation of successive differences); ***rmssd*** (root mean square of successive differences); ***pnn20*** (proportion of normal to normal > 20 ms: how many elements of *RR*_*diff*_ are larger than 20 ms); ***pnn50***; ***tpr***_*diff*_ (tpr computed on *RR*_*diff*_).

#### SDPPG features

The second derivative of the PPG processed signal (SDPPG) has been linked to chronological age^16^. During each heart beat cycle the 5 typical points (***a*** - ***e***) can be identified (Supplementary Fig. S1). To identify these points we computed the FDPPG (fourth derivative of PPG) for locating the zero crossings, which correspond to the inflection points in the SDPPG. Once two consecutive inflection points were found, only one local extreme can exist between them. The first and highest maximum of a beat cycle can be determined as ***a***, and the subsequent points, (***b***, ***c***, ***d*** and ***e***) can be detected as next minima or maxima starting from the previous one (Supplementary Fig. S1). After identifying these points, the features obtained from SDPPG are given by the amplitudes, the slopes, and the time distances.

The list of 38 features extracted based on RR, RR_**diff**_, and SDPPG is given in Supplementary Materials (Section 3 and Supplementary Fig. S4).

#### Quality thresholding

A quality score (*Q*) of the signal is computed by taking into account bad demodulation (through the variance of detected peaks height) and noise (expressed as a number of local extrema that are not detected as peaks or their corresponding valleys):

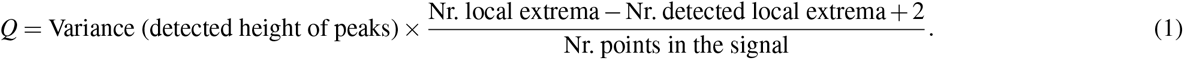

Note that +2 is used to avoid negative numbers in case of a detected peak having no valley before or after itself due to the length of its signal. The *Q* score indicates how noisy and badly demodulated the signal is; the higher the score the poorer the quality, such that 0 is the perfect score. The threshold value of *Q* was set to 0.01. By applying quality thresholding after the feature extraction phase, a flexible choice of *Q* is possible without repeating the feature extraction step.

### Prediction and classification of HVA

After discarding incomplete data (e.g. missing age label) and quality thresholding with *Q* < 0.01, the cleaned database consists of 3612 subjects: 2205 males and 1407 females, with an average age of 49 years and a standard deviation of 14.5 years. Next, the “continuous” outcome, age, was dichotomized into two classes: young (18 - 38 years, a proxy for HVA) coded 1 and old (60 - 79 years, non-HVA) coded 0. By partitioning the data into a train (2709 subjects) and a test (903 subjects) set, we train and validate the model (both the ML and DL approaches) on a training set (Fig. 1). The final model (ML) and hyperparameters (DL) were validated using the held-out test set. The age distribution of the training and test set was kept similar to that of the whole database.

#### Ridge regression (ML)

Prior to the analysis, 38 extracted features were robustly standardized by subtracting the median and dividing by the interquartile range. Since some features were highly correlated (dark red or blue colors in Fig. 3) and there may be mutlicollinearity issues, a penalized regression can be considered. Shrinking the coefficient values towards zero in multiple regressions allows the less contributing variables to have a coefficient close or equal to zero. To jointly select the relevant features for HVA, we applied ridge penalized regression; linear and logistic ridge regressions were employed for continuous and binary age outcomes, respectively^21,22^. To obtain the final model, 2/3 of the training dataset were randomly sampled and used to fit the model; for fine-tuning of the penalty parameters a 10-fold cross-validation (CV) was applied. This procedure was repeated 100 times for both continuous age (using linear ridge) and for dichotomized age (using logistic ridge). The resulting rankings of the coefficients were recorded, and averaged over 100 runs. The final score for the ranking was achieved by summing up the two scores. Then we selected some combinations of the most relevant features and we evaluated their HVA/non-HVA classification performance using Support Vector Machine (SVM) classifier (previously tuned with a 3-fold CV on training set) on the test set.

**Figure 3.**
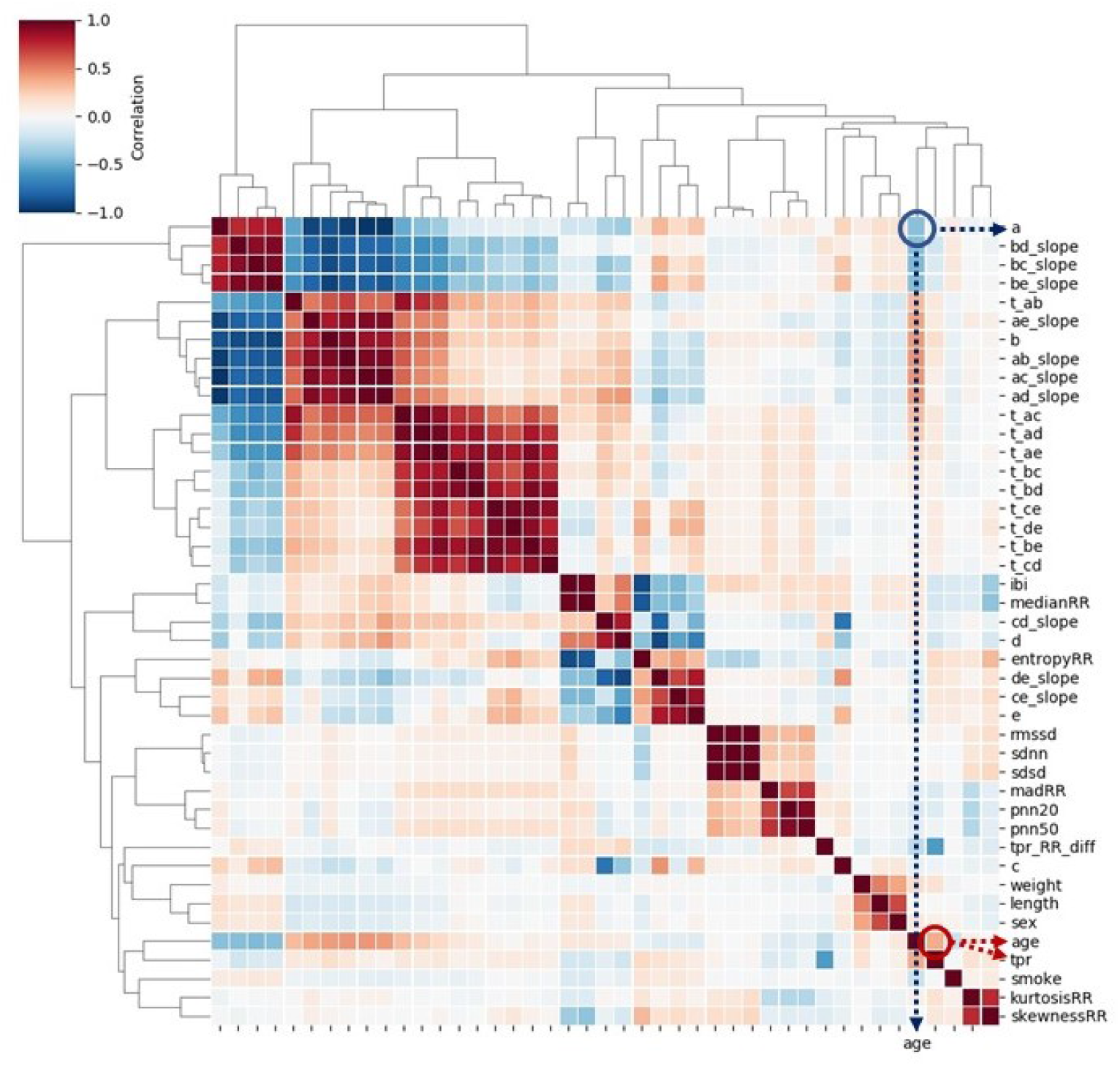
Heatmap of Pearson correlation between 38 features extracted from PPG, four covariates, and age: the feature ***a*** and age shows negative correlation, while ***tpr*** and age are positively correlated.

#### Convolutional Neural Networks (DL)

For convolutional neural networks (CNNs), the processed signals (not the extracted features) were used as input, and the dichotomized age as target. Among many CNNs, ResNet, DenseNet and WaveNet were considered. The first two have the merit of avoiding many gradient vanishing problems; they feed directly any convolutional layer using the sum (ResNet) and the concatenation (DenseNet) of all the previous layers’ output, respectively. Moreover, ResNet was found to be highly effective with some time series datasets^23^, while DenseNet used on PPG signals has already shown good results^24^. Alternatively, WaveNet was proposed to accommodate the specific nature of time sequences^25^, by using a different kind of time-oriented convolution, which is basically a truncated dilated convolution. CNNs were trained, epoch by epoch, using 2/3 of the training dataset. Their performance was validated using the remaining 1/3 of the training set; for each hyperparameter, many different values were tried out and the best was determined.

#### Evaluation of the performance of ML and DL approaches

The optimized models by ML and DL were validated on the held-out test set. Prediction performance of five models were compared using AUC: four covariates (weight, height, sex, smoking); the best two PPG features (***a*** and ***tpr***) based on ML; covariates and these two features; the best performing CNN; covariates and 38 PPG features.

## Results

The workflow is depicted in Figure 1, and the details are described in the methods section. The extracted features from PPG are given in Supplementary Materials (Section 3). All the algorithms and analysis have been performed using Python 3.7 and its suitable Anaconda distribution^26^. The code for preprocessing, feature extraction, ML and DL analysis can be found on GitHub, https://github.com/Nico-Curti/cardio, and https://github.com/LorenzoDallOlio/vascular-ageing.

### Exploratory data analysis

To visualize how strong the correlations are among age, four covariates (sex, weight, height, and smoking) and 38 PPG features (extracted from the PPG signals), a heatmap based on Pearson correlation is given in Figure 3. The strongest correlation between age and PPG features was found in ***a*** and ***tpr*** (*r* ≥ 0.35). We further investigated how well the classifier for healthy vascular aging (HVA) would perform using all extracted PPG features, spectral embedding^26^ for non-linear dimensionality reduction is first employed to cluster similar features. Figure 4 shows a relatively good separation between biologically young (blue points, indicating HVA) and old (red points, indicating non-HVA), which shows a gradual transition from HVA to non-HVA.

**Figure 4.**
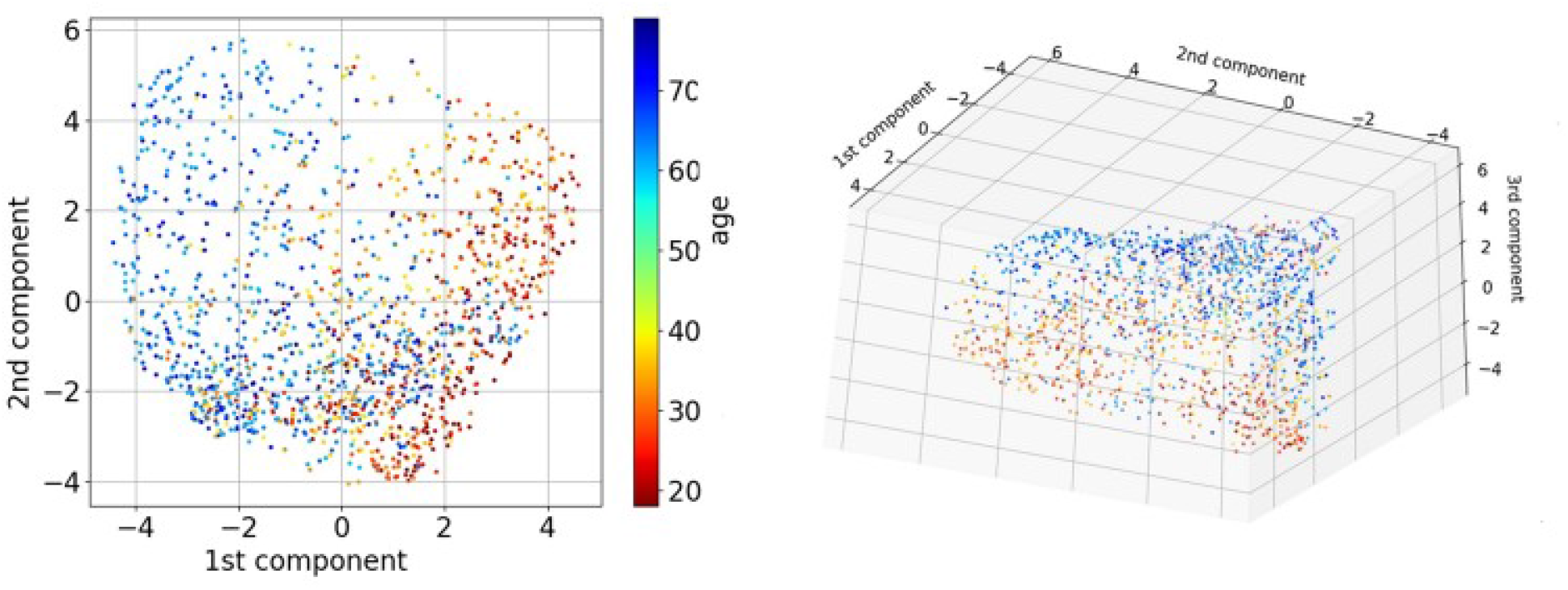
Dimension reduction of 38 PPG features extracted by spectral embedding: the left depicts the first two components, while the right depicts the first three components.

### Application of ML and DL to predict HVA

The final scores achieved by linear and logistic ridge regression are given in Table 1. The best performing PPG features were ***a*** (from SDPPG) and ***tpr*** (from PPG). Instead of just extracting features from PPG or SDPPG signals and then applying ML to these features, we also considered several CNNs using the raw signals as input. Application of CNN is *heavy*, regarding the computational burden, and tuning hyperparameters (to train CNN) is difficult as there are numerous parameters to configure. In Table 2 we report the values of hyperparameters used for WaveNet, DenseNet, and ResNet. The bold-faced ones show the hyperparameters of the best model based on the validation set performances.

**Table 1.** Results from ML approach: top six ranked variables from four covariates (smoking, sex, weight, and height) and 38 features extracted from PPG. The scores of linear and logistic ridge were achieved by ranking the absolute value of regression coefficients in decreasing order and then those rankings across 100 repetitions. The final score shows the sum of linear and logistic average rankings.

| Variable name | Linear ridge | Logistic ridge | Final ranking |
| --- | --- | --- | --- |
| smoking | 1.89 | 1.76 | 3.65 |
| sex | 4.81 | 4.67 | 9.48 |
| a | 6.64 | 10.06 | 16.70 |
| tpr | 10.61 | 7.56 | 18.17 |
| ibi | 12.36 | 8.81 | 21.17 |
| pnn20 | 14.72 | 7.65 | 22.37 |

**Table 2.** Results from DL approach: different values of hyperparameters for CNNs (WaveNet, DenseNet, and ResNet). The hyperparameter patience indicates the number of epochs to wait before early stop if no improvement in the loss function is achieved. There is no optimal dilation rate value since it is only used for WaveNets, while the best model was a ResNet CNN.

| Hyperparameter | Compared options |
| --- | --- |
| kind of CNN | WaveNet, DenseNet, **ResNet** |
| Hidden layers number | 4 8 **12** |
| dilation rate (only for WaveNets) | 1 2 4 8 |
| Batch dimension | 4 16 **32** 64 256 |
| Kernel dimension | 3 6 10 30 **50** 80 |
| Activation function | ReLU **ELU** SELU |
| Optimizer | SGD ADAM **NADAM** |
| Start learning rate | $10^{-1}$ $10^{-2}$ $10^{-3}$ **$10^{-4}$** $10^{-5}$ |
| Patience | 5 **8** 10 |

### Evaluation of prediction performance

To validate the results, the independent test set was used (Fig. 1). The AUC obtained using only one feature (i.e., ***a***, ***slope-AC***, or ***tpr***) was around 0.8 (Supplementary Table S1). The following models are considered: (i) covariates (weight, height, sex, smoking), (ii) the best two PPG features (***a*** and ***tpr***) from ML, (iii) covariates and these two features, (iv) the best performing CNN, and (v) covariates and all PPG features. One should take note that the model (v) is not recommended due to the overfitting, which may not lead to such a good performance on a different dataset. For comparing the prediction performance of the five models, the area under the ROC curve (AUC) was computed, and the results are depicted in Figure 5. By adding the PPG features (***tpr*** and ***a***) to the covariates, AUC increased from 0.742 to 0.947. The 12-layer ResNet model (AUC=0.953) performed similar to including all variables (AUC=0.954).

**Figure 5.**
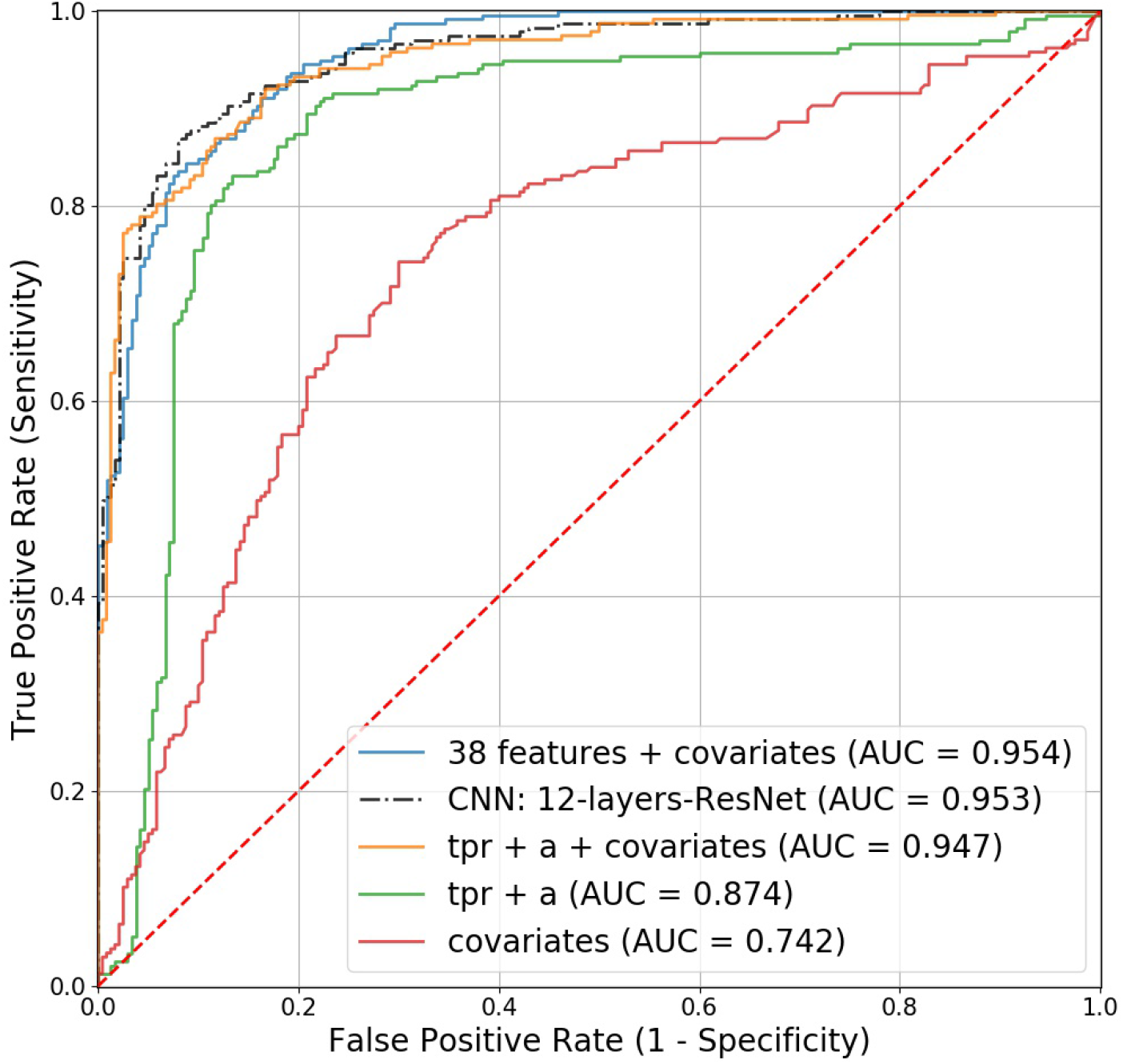
Receiver operating characteristic (ROC) curves for the competing models in predicting healthy vascular aging (HVA): the four covariates include sex, weight, height, and smoking.

### Sex-stratified analysis

There are differences in body weight, height, body fat distribution, heart rate, stroke volume, and arterial compliance between the two sexes. In the very elderly, age related large artery stiffness is reported to be more pronounced in women^11^. Therefore, based on the two features obtained by the best performing ML, we investigated whether men and women age differently regarding HVA. The model, age = ***a*** + ***tpr*** + covariates + error, was used to estimate coefficients using the training set. The obtained coefficient estimates were used to predict vascular aging using the test set. The results are shown in Figure 6. The points in the figures are the predicted age of each male (blue) and female (red). To depict the fitted trend line the locally weighted scatterplot smoothing (LOWESS) technique is used. For clear comparison, two fitted lines are presented (in right panel). In average, females are healthier than males regarding vascular aging. Focusing on the individual feature, the ***a*** wave from SDPPG showed the considerable difference between men and women (Supplementary Fig. S5).

**Figure 6.**
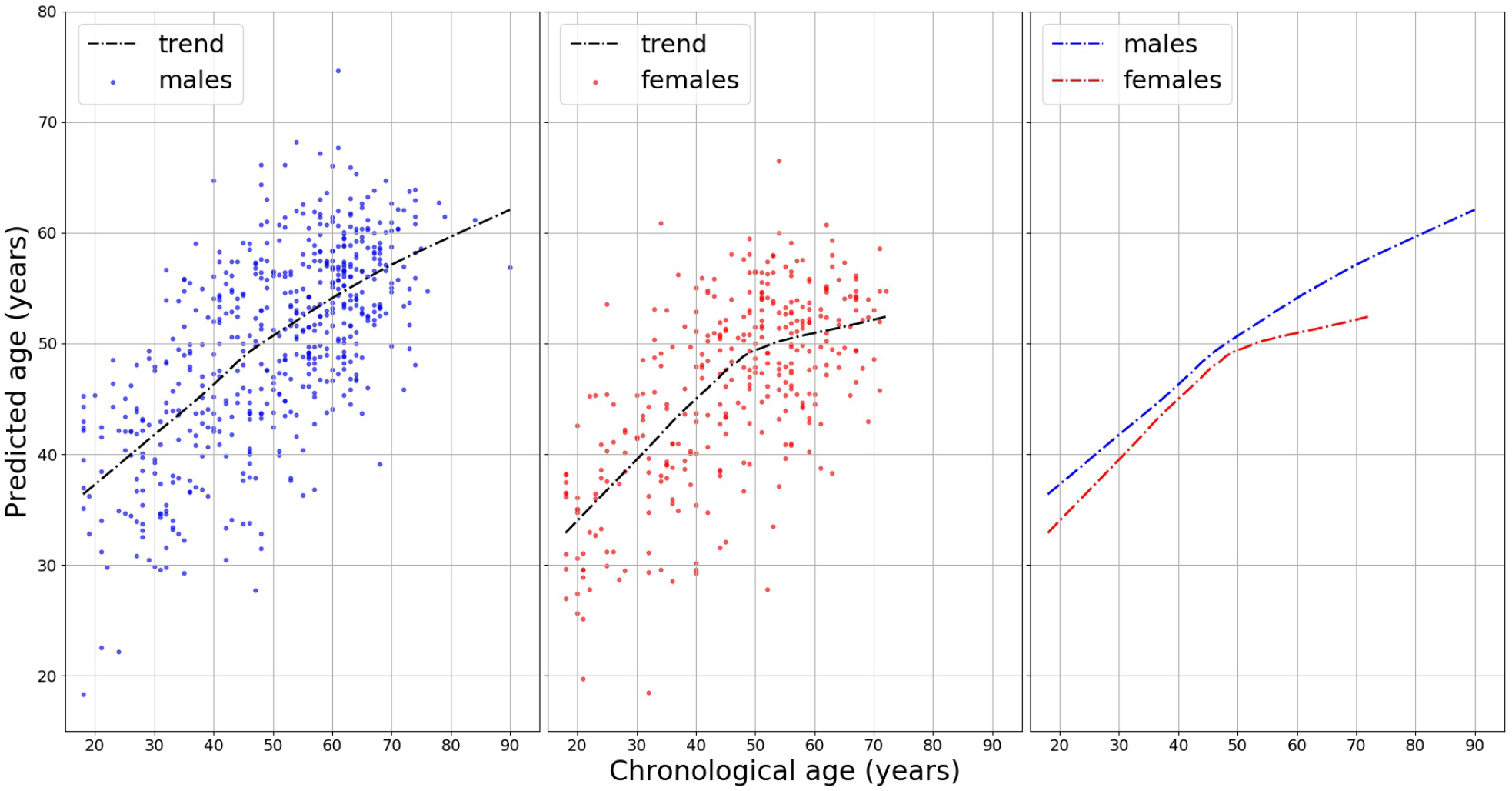
Sex-stratified analysis for predicting vascular age: a locally weighted scatterplot smoothing (LOWESS) of the ages predicted by the model ***a***+***tpr***+covariates for men and women, separately. The figure in right panel shows that average predicted age for females is always lower than that for males, indicating females have healthier vascular aging. Moreover, for females, a reduction in the trend slope around 50 years old is shown (in middle panel).

## Discussion

Our work has been motivated by the unique crowd-sourced data from Heart for Heart initiative. Aging is the main risk factor for vascular disease. The concept of biological aging comes from the fact that individuals do not age at the same pace. In particular, vascular aging involves arterial degeneration and hardening that impairs vascular function and ultimately causes end organ damage^27^. Concerning biomarkers of biological vascular aging, the most common arterial stiffness measure is pulse wave velocity (PWV), in our case the PPG waves, the velocity at which the blood pressure wave moves along the arterial tree. From the statistical point of view, biological aging can be studied in two ways. Firstly, one can study individual differences as in the brain aging process to reflect advanced or delayed brain ageing.^28^. Typically a linear regression is employed to show the differences in slope (see Figure 1 in Hamczyk *et al*.^27^), or to compare two age groups using Pearson correlation coefficients^29^. Secondly, as in the research of healthy aging and longevity^30,31^, one can consider two groups of *young* and *old*. Through a classification task, one can estimate the probability of belonging to either class and determine suitable biomarkers for longevity potential. Our approach here is the second one, which can be extended to monitor other diseases such as Atrial Fibrillation. Also, publicly available PPG data that includes ‘raw’ signals are still rare. Having abundant PPG data from the general population, we aimed to scrutinize the whole process from data preprocessing of the PPG signals to the prediction of HVA using PPG.

Recently deep neural network approaches have grown in popularity^24,32^, there are several challenges applying DCNN (deep convolutional neural network) to health data. One needs to choose the right topology of NN: how many hidden layers, how do you trade off the number of parameters versus the amount of training data, etc. Computational power is one of the biggest limitation, when the aim is to get instant diagnosis using a wearable device^33^. Moreover, the algorithms deployed are inscrutable (it is a black-box approach) and it is difficult to interpret their results compared to other ML approaches. Prior to employing ML, several steps are involved: the signals have to be detrended and demodulated, the quality of the signals has to be assessed and signals of poor quality should be removed, peak detection algorithm has to be applied, relevant features must be extracted. Instead of using some aspects extracted from the signal, the whole signal is used as input for DL; the latter might be advantageous, when certain hidden features were not included in the model for ML. Therefore, we have investigated effectiveness of applying the computer intensive DL methods (ResNet, DenseNet, WaveNet) against the relatively simple ML method (ridge penalized regression).

The prediction performance indicated the best DL (AUC of 95.3%), 12-layers ResNet, slightly outperformed the ML (AUC, sensitivity, and specificity of 94.7%, 87.3%, and 86.3%) with the model, 2 PPG features (***a*** and ***tpr***) + 4 covariates (sex, weight, height, smoking). Nevertheless, ML had the merit of identifying potential biomarkers. It has been reported that features derived from the contour of PPG signals showed association with age, in particular regarding arterial elasticity and the changes in the elastic properties of the vascular system^15^. The feature ***a*** wave extracted from SDPPG and ***tpr*** from PPG showed negative and positive correlation with age, respectively. The SDPPG features derived from the amplitudes of the distinctive waves situated in the systolic phase of the heart cycle, quantify the acceleration of the arterial blood vessels’ walls. ***tpr*** (turning point ratio) is based on the non-parametric ‘Runs Test’ to measure the randomness in a time-series, and it is higher when series are more random: for instance, AF patients tend to have higher ***tpr***^20^. Taking age as a proxy for vascular aging, both larger ***a*** and lower ***tpr*** predict healthy vascular aging (HVA). In addition vascular aging differs between the sexes. Age-related changes in vascular function generally include increasing endothelial dysfunction and arterial stiffness^34^. We have shown that women appeared to be healthier (younger) than men regarding vascular aging (Fig. 6). Moreover, the clear slope variation in the females of Figure 6 might be interpreted as slowing down the vascular aging process, ultimately leading to longer life expectancy of women. The best performing ML model (***a*** + ***tpr*** + sex + weight + height + smoking) supports these findings (Fig. 6). For predicting HVA, therefore, ML using a few PPG features together with relevant clinical information can be a viable option against computer-intensive DL approaches^33^. Further, this study might be considered as a proof of concept for prediction of vascular aging based on chronological age. Extending and fine-tuning of the best performing ML model may lead to a risk score for vascular aging. Adding other relevant variables (the known risk factors) into the current model can be an option. Taking PPG measurements by smartphone camera together with simple algorithm for feature extraction and prediction for vascular aging facilitates self-monitoring of individual risk score, which can be linked to smoking and eating habits.

Note that CNNs investigated here demonstrated good performances obtained by simple structures (never more than 12 hidden layers, which is feasible for common laptop to train). When one’s interest is studying aging process^27^, for demonstration purpose, we slightly modified our approach as reported in Supplementary Materials (Section 4 and Supplementary Fig. S6–S8). Further fine-tuning of CNN to predict a continuous outcome is needed^35^, but beyond the scope of this work. When accurate prediction is the main purpose and any distinct features need to be fully incorporated (as for AF detection), CNNs can be good candidates. Our results with age as a target can be employed for transfer learning approaches for a specific target such as AF.

We showed that PPG measured by smartphone has the potential for large scale, non-invasive, patient-led screening. However, current evidence is often biased due to low quality studies, black-box methods, or small sample sizes. For instance, the reliability of ultra-short HRV features (PPG measurements less than 5 minutes) remains unclear and many HRV analyses have been conducted without questioning their validity^36^. On the other end of spectrum, systematic review of assessing the balance of benefits and harms of screening for AF with smartphone acquired PPG is lacking. This work contributes to establishing generally accepted algorithm based on open data and software, which is of major importance to reproduce the procedures, and to further improve and develop methods.

## Acknowledgements

This work has received support from the EU/EFPIA Innovative Medicines Initiative 2 Joint Undertaking BigData@Heart grant (116074), and from the European Union’s Horizon 2020 research and innovation programme IMforFUTURE under H2020-MSCA-ITN grant agreement number 721815. Data for this study come from the Heart for Heart (H4H) initiative. Permission was obtained to use data for this study, and Happitech has shared research data. F.W.A. is supported by UCL Hospitals NIHR Biomedical Research Centre.

## Author contributions statement

H.W.U., G.C., and F.W.A. conceived and designed the study. L.D. and H.W.U. analyzed and interpreted the data, and drafted the manuscript. N.C. and D.R. helped with the initial part of the code for the PPG data preprocessing. All authors reviewed the manuscript.

## Additional information

### Supplementary Information

Supplementary Materials.

## References

1. Matsumura, K., Rolfe, P. & Yamakoshi, T. iPhysioMeter: a smartphone photoplethysmograph for measuring various physiological indices. Methods Mol. Biol. 1256, 305–326, DOI: 10.1007/978-1-4939-2172-0_21 (2015).

2. Krivoshei, L. et al. Smart detection of atrial fibrillation. Europace 19, 753–757, DOI: 10.1093/europace/euw125 (2017).

3. Castaneda, D., Esparza, A., Ghamari, M., Soltanpur, C. & Nazeran, H. A review on wearable photoplethysmography sensors and their potential future applications in health care. Int. J. Biosens. & Bioelectron. 4, 195–202, DOI: 10.15406/ijbsbe.2018.04.00125 (2018).

4. Li, K. H. C. et al. The current state of mobile phone apps for monitoring heart rate, heart rate variability, and atrial fibrillation: narrative review. JMIR Mhealth Uhealth 15, e11606, DOI: 10.2196/11606 (2019).

5. Chan, P. H. et al. Diagnostic performance of a smartphone-based photoplethysmographic application for atrial fibrillation screening in a primary care setting. J. Am. Hear. Assoc. 5, 27444506, DOI: 10.1161/JAHA.116.003428 (2016).

6. De Ridder, B., Van Rompaey, B., Kampen, J. K., Haine, S. & Dilles, T. Smartphone apps using photoplethysmography for heart rate monitoring: meta-analysis. JMIR Cardio 2, e4, DOI: 10.2196/cardio.8802 (2018).

7. Jonas, D. E. et al. Screening for atrial fibrillation with electrocardiography: an evidence review for the u.s. preventive services task force. JAMA 320, 485–498, DOI: 10.1001/jama.2018.419 (2018).

8. Sudler & Hennessey. Heart For Heart. Website http://www.heartrateapp.com/ (2020).

9. Happitech. Monitor your heart rhythm using only a smartphone. Smartphone App http://www.happitech.com (2020).

10. Ghebre, Y. T., Yakubov, E., Wong, W. T. & Krishnamurthy, P. Vascular aging: implications for cardiovascular disease and therapy. Transl. Medicine 06, 183, DOI: 10.4172/2161-1025.1000183 (2016).

11. Jani, B. & Rajkumar, C. Ageing and vascular ageing. Postgrad. Med. J. 82, 357–362, DOI: 10.1136/pgmj.2005.036053 (2006).

12. North, B. J. & Sinclair, D. A. The intersection between aging and cardiovascular disease. Circ. Res. 110, 1097–1108, DOI: 10.1161/CIRCRESAHA.111.246876 (2012).

13. Laina, A., Stellos, K. & Stamatelopoulos, K. Vascular ageing: underlying mechanisms and clinical implications. Exp. gerontology 109, 16–30, DOI: 10.1016/j.exger.2017.06.007 (2018).

14. Nilsson, P. M. et al. Characteristics of healthy vascular ageing in pooled population-based cohort studies: the global metabolic syndrome and artery research consortium. J. Hypertens. 36, 2340–2349, DOI: 10.1097/HJH.0000000000001824 (2018).

15. Yousef, Q., Reaz, M. B. & Ali, M. A. The analysis of PPG morphology: investigating the effects of aging on arterial compliance. Meas. Sci. Rev. 12, 266–271, DOI: 10.2478/v10048-012-0036-3 (2012).

16. Pilt, K. et al. New photoplethysmographic signal analysis algorithm for arterial stiffness estimation. The Sci. World J. 169035, DOI: 10.1155/2013/169035 (2013).

17. Ahn, J. M. New aging index using signal features of both photoplethysmograms and acceleration plethysmograms. Healthc. Informatics Res. 23, 53–59, DOI: 10.4258/hir.2017.23.1.53 (2017).

18. Tamura, T., Maeda, Y., Sekine, M. & Yoshida, M. Wearable photoplethysmographic sensors—past and present. Electronics 3, 282–302, DOI: 10.3390/electronics3020282 (2014).

19. van Gent, P., Farah, H., van Nes, N. & van Arem, B. HeartPy: HeartPy: a novel heart rate algorithm for the analysis of noisy signals. Transp. Res. Part F: Traffic Psychol. Behav. 66, 368–378, DOI: 10.1016/j.trf.2019.09.015 (2019).

20. Tang, S.-C. et al. Identification of atrial fibrillation by quantitative analyses of fingertip photoplethysmogram. Sci. Reports 7, 45644, DOI: 10.1038/srep45644 (2017).

21. Hoerl, A. E. & Kennard, R. W. Ridge regression: biased estimation for nonorthogonal problems. Technometrics 12, 55–67 (1970).

22. Hastie, T., Tibshirani, R. & Friedman, J. H. J. H. The elements of statistical learning : data mining, inference, and prediction (Springer, 2001).

23. Ismail Fawaz, H., Forestier, G., Weber, J., Idoumghar, L. & Muller, P.-A. Deep learning for time series classification: a review, DOI: 10.1007/s10618-019-00619-1 (2019).

24. Poh, M.-Z. et al. Diagnostic assessment of a deep learning system for detecting atrial fibrillation in pulse waveforms. Heart 104, 1921–1928, DOI: 10.1136/heartjnl-2018-313147 (2018).

25. van den Oord, A. et al. WaveNet: a generative model for raw audio. Arxiv http://dx.doi.org/10.6084/m9.figshare.853801 (2016).

26. Anaconda. Anaconda Software Distribution. Software http://www.anaconda.com (2016).

27. Hamczyk, M. R., Nevado, R. M., Barettino, A., Fuster, V. & Andrés, V. Biological Versus Chronological Aging: JACC Focus Seminar. J. Am. Coll. Cardiol. 75, 919–930, DOI: 10.1016/j.jacc.2019.11.062 (2020).

28. Cole, J. H. & Franke, K. Predicting Age Using Neuroimaging: Innovative Brain Ageing Biomarkers. Trends Neurosci. 40, 681–690, DOI: 10.1016/j.tins.2017.10.001 (2017).

29. Remondini, D. et al. Identification of a T cell gene expression clock obtained by exploiting a MZ twin design. Sci. Reports 7, 1–8, DOI: 10.1038/s41598-017-05694-2 (2017).

30. Gonzalez-Covarrubias, V. et al. Lipidomics of familial longevity. Aging Cell 12, 426–434, DOI: 10.1111/acel.12064 (2013).

31. Beekman, M. et al. Classification for longevity potential: The use of novel biomarkers. Front. Public Heal. 4, 28, DOI: 10.3389/FPUBH.2016.00233 (2016).

32. Kwon, S. et al. Deep learning approaches to detect atrial fibrillation using photoplethysmographic signals: algorithms development study. JMIR Mhealth Uhealth 21, e12770, DOI: 10.2196/12770 (2019).

33. Bashar, S. K. et al. Atrial fibrillation detection from wrist photoplethysmography signals using smartwatches. Sci. Reports 9, 15054, DOI: 10.1038/s41598-019-49092-2 (2019).

34. Merz, A. A. & Cheng, S. Sex differences in cardiovascular ageing. Heart 102, 825–831, DOI: 10.1136/heartjnl-2015-308769 (2016).

35. Biswas, D. et al. CorNET: Deep Learning Framework for PPG-Based Heart Rate Estimation and Biometric Identification in Ambulant Environment. IEEE Transactions on Biomed. Circuits Syst. 13, DOI: 10.1109/TBCAS.2019.2892297 (2019).

36. Pecchia, L., Castaldo, R., Montesinos, L. & Melillo, P. Are ultra-short heart rate variability features good surrogates of short-term ones? State-of-the-art review and recommendations. Healthc. Technol. Lett. 5, 94–100, DOI: 10.1049/htl.2017.0090 (2018).

