## Supplementary_PPG_vascular_Ageing for "Prediction of vascular aging based on smartphone acquired PPG signals" (from another version of this preprint: version 1, 2020-05-28)

**Title: Prediction of vascular ageing based on smartphone acquired PPG signals**

**Supplementary Materials**

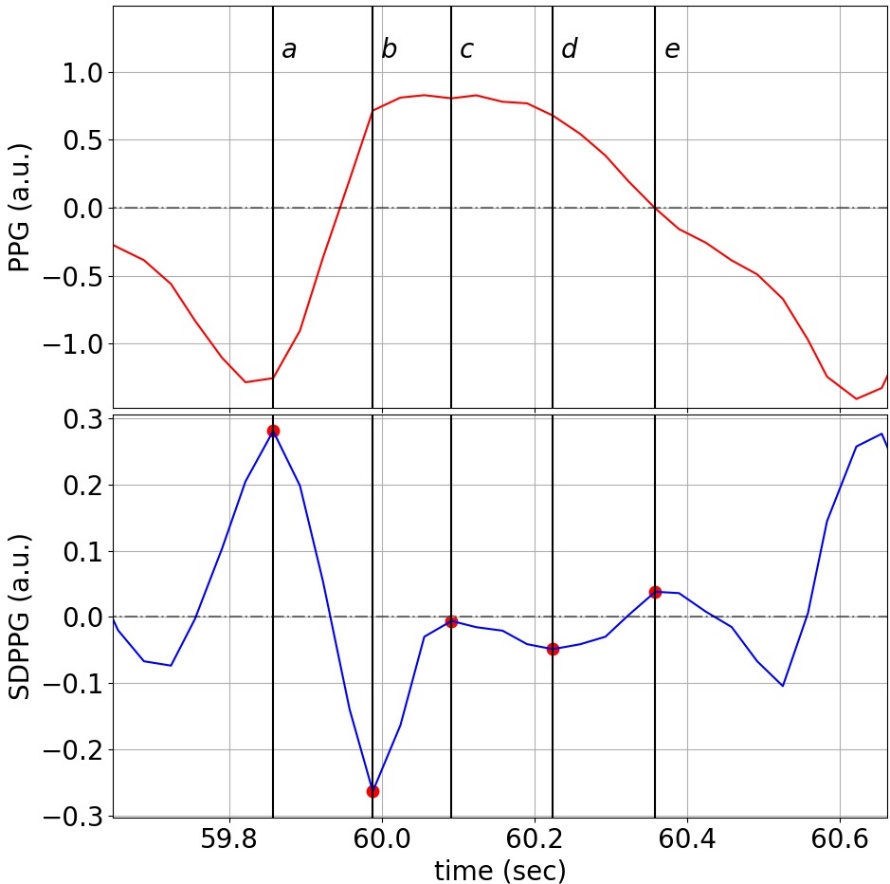

**Figure S1:** PPG (top) and the corresponding SDPPG (bottom) shapes. Red points represent the peaks and changing points, ***a, b, c, d,*** and ***e*** identified by SDPPG.

**1. The list of 38 features extracted from the PPG signals**

i. SDPPG group:
   - ***a***, ***b***, ***c***, ***d***, ***e*** (See Supplementary Fig. S1), ***ab_slope*** (slope of the segment connecting point ***a*** and point ***b*** in the SDPPG), ***ac_slope***, ***ad_slope***, ***ae_slope***, ***bc_slope***, ***bd_slope***, ***be_slope***, ***cd_slope***, ***ce_slope***, ***de_slope***, ***t_ab*** (time interval that separates point ***a*** and point ***b*** in the SDPPG), ***t_ac***, ***t_ad***, ***t_ae***, ***t_bc***, ***t_bd***, ***t_be***, ***t_cd***, ***t_ce***, ***t_de***.
ii. PPG group:
    - Extracted from RR: ***ibi*** (average inter-beat interval - the mean of RR), ***medianRR***, ***entropyRR***, ***kurtosisRR***, ***skewnessRR***, ***madRR*** (median absolute distance of RR), ***tpr*** (turning point ratio of RR - the number of local extrema in RR divided by the number of elements in RR), ***sdnn*** (standard deviation normal to normal, basically standard deviation of RR).
    - Extracted from *RR_di_ _f_ _f_* : ***pnn20*** (proportion of normal to normal 20, the percentage of elements of RR which differ of more than 20 ms from the successive element - the percentage of elements of *RR_di_ _f_ _f_* greater than 20 ms), ***pnn50***, ***sdsd*** (standard deviation of successive differences), ***rmssd*** (root mean square of successive differences), ***tpr_RR_diff.***

**Table S1.** Sensitivity, Specificity and AUC for various models. Included four covariates are: sex, weight, height, and smoking.

| **Model** | **Sensitivity (%)** | **Specificity (%)** | **AUC (%)** |
| --- | --- | --- | --- |
| 38 PPG features + 4 covariates | 87.76 | 85.42 | 95.43 |
| CNN: 12-layers-ResNet | 90.72 | 85.83 | 95.34 |
| ***a*** + ***tpr*** + 4 covariate | 87.34 | 86.25 | 94.73 |
| ***ac_slope*** + ***tpr*** + 4 covariates | 87.76 | 88.33 | 94.60 |
| *a* + ***tpr*** | 83.97 | 82.50 | 87.43 |
| ***ac_slope*** | 78.06 | 78.75 | 81.68 |
| ***a*** | 77.64 | 78.33 | 80.79 |
| ***tpr*** | 73.00 | 76.25 | 80.50 |
| 4 covariates | 69.62 | 72.50 | 74.21 |
| ***pnn20*** | 66.67 | 61.67 | 66.29 |
| ***ibi*** | 40.93 | 72.50 | 55.21 |

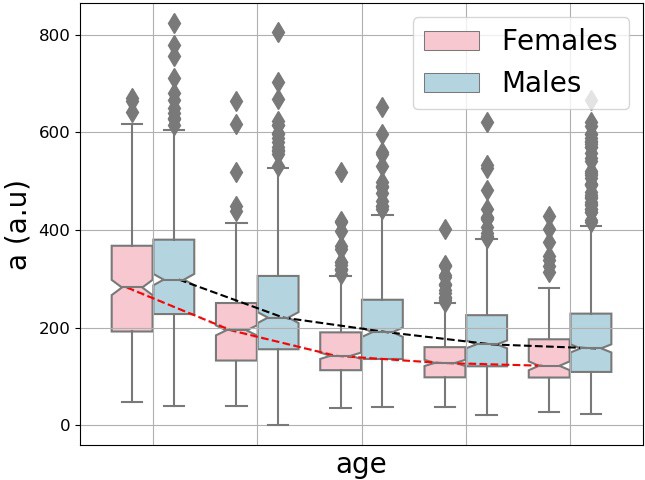

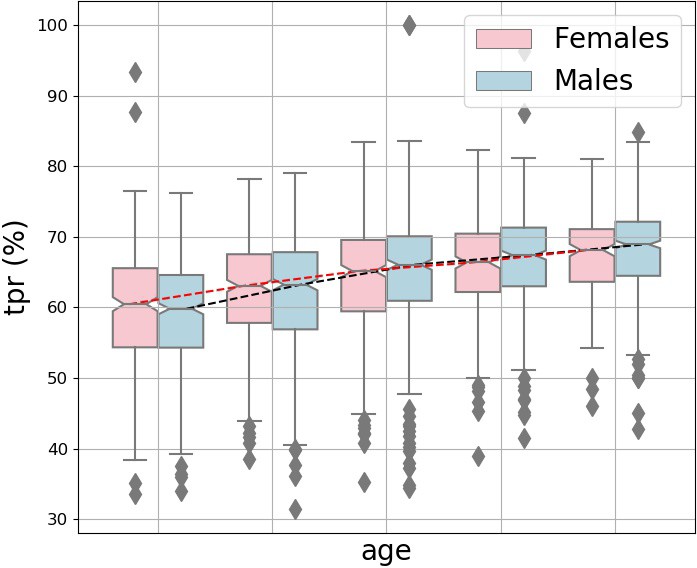

**Figure S2:** Sex-stratified analysis. The chronological age was categorized into quintiles: (17, 35], (35, 47], (47, 55], (55, 62], (62, 98]. Left: the feature ***a*** shows negative correlation with age, and females having lower ***a*** than males in the same age category. Right: the feature ***tpr*** is positively correlated with age, and shows little differences between males and females.
